# Enhanced mouse virulence of mpox virus clade Ib over clade IIb despite genomic changes caused by human-to-human transmission

**DOI:** 10.1101/2025.09.22.677787

**Authors:** Alazne R. Unanue, Rocío Martín, Carolina Sánchez, Ana Moraga-Quintanilla, Laura F. del Ama, Pedro J. Sánchez-Cordón, Antonio Alcamí, Bruno Hernáez

**Affiliations:** Centro de Biología Molecular Severo Ochoa, Consejo Superior de Investigaciones Científicas (CSIC)-Universidad Autónoma de Madrid (UAM), 28049, Madrid, Spain; Centro de Investigación en Sanidad Animal (CISA), Instituto Nacional de Investigación y Tecnología Agraria y Alimentaria (INIA), Consejo Superior de Investigaciones Científicas (CSIC), Valdeolmos, 28130, Madrid, Spain

**Author notes:** Both authors equally contributed to this work (Rocío Martín, Carolina Sánchez). Author order was determined by alphabetical order.

**Keywords:** mpox, monkeypox virus, clade 1b, human-to-human transmission, APOBEC3, virulence

## Abstract

The recently emerged clade Ib of mpox virus (MPXV) is spreading rapidly across Central and West Africa raising concerns about its potential virulence. Similar to clade IIb lineage B.1, which was responsible for the 2022 global outbreak, clade Ib exhibits sustained human-to-human transmission and a pattern of APOBEC3-associated genomic mutations. Here, we show that clade Ib displays enhanced cell-to-cell dissemination *in vitro* compared to clade IIb. Additionally, using the CAST mouse model, we show that clade Ib retains a higher level of virulence than that of the markedly attenuated clade IIb. Clade Ib leads to significant weight loss and high mortality in animals following both intraperitoneal and intranasal challenge. Histopathological analysis revealed more severe and extensive lung lesions in clade Ib–infected animals, accompanied by a broader distribution of viral antigens. Moreover, clade Ib, unlike IIb, disseminated efficiently to internal organs. These findings indicate that clade Ib MPXV has not undergone attenuation after human-to-human transmission to the extent observed in clade IIb and underscore the need for surveillance and preparedness against new emerging MPXV lineages.

## Introduction

Human mpox, formerly known as monkeypox, has been historically considered as a zoonotic viral disease since mpox virus (MPXV) is sporadically transmitted from rodents or nonhuman primates in Central and West Africa to humans ^1,2^. MPXV belongs to the Orthopoxvirus genus of the family *Poxviridae* and two genetically distinct clades have been identified: clades I and II, originally designed as Central and Western Africa, respectively ^3,4^. The clade I causes a more severe form of the disease, which resembles a milder form of smallpox with characteristic skin lesions, fever, swollen lymph nodes and a case fatality rate around 10% in endemic regions. In contrast, case fatality rate associated to clade II is below 2%. Recently, clade II has been further classified into subclade IIa and IIb ^5^. Mpox was first observed in 1970 in the Democratic Republic of the Congo (DRC) ^6^, the sole country continually reporting mpox cases in the last five decades. Since then, MPXV has caused diverse outbreaks out of endemic regions, always originated by imported cases of clade II viruses with limited human-to-human transmission, such as the outbreaks of 2003 in the USA and 2018 in the UK caused by the importation of rodents from endemic areas and a traveller from Nigeria, respectively ^7,8^.

However, in 2022 the epidemiological scenario changed completely, as World Health Organization (WHO) declared a Public Health of International Concern for mpox. During the year that this emergency remained active, more than 94,000 human cases were confirmed in 130 countries, with 157 deaths. This global outbreak was caused by the emerging clade IIb lineage B.1 ^9,10^, characterized by an unprecedent human-to-human transmission rate, with 98% of cases among young men, most of them identified as men who have sex with men. Genomic analyses revealed that most of the nucleotide differences in the emerged global lineage B.1, compared to its predecessor, are consistent with apolipoprotein B mRNA editing enzyme catalytic polypeptide 3 (APOBEC3)-mediated editing as a result of sustained human transmission, as has been described for other MPXV lineages and other orthopoxviruses infecting humans ^11,12^. Since 2023 the number of confirmed mpox cases caused by clade IIb viruses decreased and it is still reported at low levels in many countries.

This disease once again became a cause for concern in 2024 due to the appearance of an emerging clade I (MPXV subclade Ib) which rapidly spread through person to person contact across Central and Eastern Africa ^13,14^, leading WHO to declare a second Public Health of International Concern ^15^. In the last year, sporadic travel-associated cases of human clade Ib infection have also been reported out of Africa, in Europe, Asia or North America ^16–19^. Preliminary observations during the clade Ib outbreak in Africa suggest that this virus may be more virulent than clade IIb exhibiting a higher case fatality rate compared to what was observed during 2022 global mpox outbreak in non-African countries. However, factors beyond clade-specific differences can confound fatality rates in African regions affected by clade Ib, such as the high proportion of people living with HIV, the concentration of cases in young children, malnutrition and weak healthcare systems, among others ^20,21^. Indeed, analysis of the MPXV genome sequencing data available from clade Ib viruses confirmed an APOBEC3 mutational signature, as previously observed in clade IIb. Additionally, similar to clade IIb viruses, the virulence gene *OPG032* encoding the viral complement control protein, is absent from clade Ib viruses while it is present in clade Ia ^22,23^. In this regard, to date, the virulence of this recently emerged clade Ib has not been addressed in any animal model of pathogenesis.

Americo et al ^24^ showed that the CAST/Eij mouse model of MPXV infection can recapitulate the pathogenicity differences among Ia, IIa and IIb B.1 MPXV clades. Despite similar replication kinetics in cell culture, the global clade IIb was shown to be highly attenuated compared to its predecessor viruses from clade IIa. Upon intranasal or intraperitoneal infection with up to 10^5^ pfu of clade IIb, no mortality or signs of disease were observed in CAST mice. In contrast, clade IIa LD_50_ value was determined around 10^3^ pfu. In this model, viral dissemination of the diverse clades to visceral organs like the spleen and liver follows the order clade I > clade IIa > clade IIb. More recently, a new study confirmed these virulence differences in CAST mice between clade IIa and IIb B.1 using intranasal and intradermal infections ^25^. Interestingly, the global clade IIb B.1 also exhibited reduced virulence compared to its closest ancestor, the clade IIb A.1, which differs by just 46 nucleotides ^9^, suggesting a relationship between the APOBEC3-mediated editing of MPXV genomes and the reduced virulence in mice ^26^. In addition, infection with clade IIb B.1 induced a higher interferon-based response than that caused by clade IIb A.1 in fibroblasts ^26^ and also in the lungs of clade IIa infected mice ^25^.

In the present work we evaluated the virulence of the recently emerged MPXV clade Ib in the CAST mice model and compared it with the global clade IIb lineage B.1, another MPXV that has also experienced high human-to-human transmission. The data presented here demonstrate that clade Ib is not as attenuated as the global clade IIb.

## Methods

### Cells and viruses

BS-C-1 cells (ATCC CCL 26), BSC-40 (ATCC CRL 2761) and human foreskin fibroblasts (HFF-1, ATCC SCRC-1041) were maintained at 37°C and 5% CO_2_ in 10% heat-inactivated fetal bovine serum (FBS) containing DMEM and supplemented with 100 U/ml penicillin, 100 μg/ml streptomycin and 2 mM L-glutamine.

MPXV Clade IIb lineage B.1 (MPXV-CBM) was isolated during the 2022 global outbreak in Madrid, Spain (Accession number GCA_964276875, ^27^ and MPXV Clade Ib (MpxV/PHAS-506/Passage-03/SWE/2024_09_11, GISAID EPI_ISL_19348512) collected in Sweden in 2024 ^17^ was sourced from European virus archive global (EVAg). Both viruses were expanded in BS-C-1 cells and purified by ultracentrifugation through a 36% (w/v) sucrose cushion prior to infect cells or mice. Viral stocks obtained were titrated twice by plaque assay in monolayers of BS-C-1 cells prior to animal infections.

### Virus growth curves

BS-C-1 cells in 12-well plates were infected for 1 h at 37 °C at high moi (5 pfu/cell) or low moi (0.01 pfu/cell) in the one-step or multi-step growth curves, respectively. Cells were then washed and fresh medium was added. Infection proceeded for 24 h in the one-step growth curve and for 72 hpi in the multi-step growth curve. For every time point, the medium was harvested and centrifuged at 1,800 × g for 5 min to pellet detached cells (extracellular virus, EV). These cells were combined with infected cells scraped from the plate into 0.5 ml of fresh medium (cell associated virus, CAV). In all cases, samples were frozen, thawed three times and titrated on BS-C-1 cells in duplicates.

### MPXV plaque assays

BS-C-1 cells seeded in 12-well plates were inoculated with 10-fold serial dilutions of the solution containing virus. After a 60 min adsorption period, cells were washed and further incubated at 37 °C in semi-solid carboxy-methyl cellulose media with 2.5% FBS. Excepted otherwise indicated, cells were fixed at 4 dpi in 10 % formaldehyde and stained with 0.1% (w/v) crystal violet.

For comet assays BSC-40 cells were seeded in 6-well plates to obtain 70% cell confluence the day of the assay. Cells were infected with 50 pfu of MPXV and after 60 min adsorption period at 37 °C cells were washed and incubated for 4 days in liquid DMEM containing 2% FBS. Viral plaques were fixed and stained as described above.

### Infection of Mice

Cast/EiJ mice breeding under pathogen-free conditions was carried out at the animal house in Centro de Biologia Molecular Severo Ochoa (CBM) in Madrid (Spain). All the experimental procedures were approved by the Ethical Review Board of CBM and CSIC, and approved by local authorities (Comunidad de Madrid) under reference PROEX 241.1/21 and performed by smallpox vaccinated investigators. Groups of 7-12 weeks (12-16 gr.) male and female Cast/EiJ mice were anesthetized by inhalation of isoflurane immediately before infection with sucrose-purified viral stocks. We and others ^25^ have not previously observed relevant differences between males and females after MPXV infection and details on the composition of the groups in the challenge experiments can be found in Supplementary information (Table S1). For intranasal infection a single 20 μl dose of PBS-0.1% BSA containing the indicated amount of MPXV-CBM or MPXV Clade Ib was directly administered on the nostril openings. For intraperitoneal infection the amount of virus was diluted in 200 ul PBS-0.1% BSA/animal. Mock infected animals were inoculated with the same volume of PBS-0.1% BSA. After infection, viral inocula used were back-titrated by plaque assay to verify the administered viral dose. Mice were housed in ventilated racks under biological safety level 3 containment facilities at CBM and monitored daily for survival, weight, temperature and signs of disease. Mouse temperature and identification data were collected by implanted radio frequency identification using transponders TP-500 and DAS-8037-IUS reader system (Bio Medic Data Systems). Animals exceeding 25% weight loss together with clear signs of disease were euthanized using carbon dioxide inhalation.

### Lung histopathology and detection of viral antigen by immunohistochemistry

Infected or mock-infected Cast/EiJ mice were euthanized at 5 dpi. The entire left lung lobes were removed from each mouse, immersion-fixed in 4% paraformaldehyde (Sigma) for 48 hours, routinely processed and embedded in paraffin wax. The paraffin blocks were used to obtain serial 4 µm sections using a rotary microtome. The sections were used for further routine histopathological studies (haematoxylin and eosin staining). Lung sections were examined microscopically using an Olympus BX43 microscope by a veterinary pathologist blinded to sample identity and group assignment. To assess the presence and severity of lung lesions, histopathological scoring parameters based on previous reports of MPXV virus infection in murine models were used ^28,29^. Histopathological examination focused on the presence of lesions in three pulmonary structures: 1) Alveoli: alveolar haemorrhages, alveolar oedema, alveolar infiltrates, thickening of alveolar septal (interstitial pneumonia), necrosis of alveolar septa/desquamation, cytopathic effect in pneumocytes or syncytia; 2) Lung vessels: vascular congestion, perivascular haemorrhages/oedema, perivascular infiltrate, vasculitis, microthrombosis; and 3) Bronchi/bronchioles: peribronchial/peribronchiolar infiltrates, epithelial hyperplasia, epithelial necrosis with detached epithelium/debris or inflammatory cells in the lumen (bronchitis/bronchiolitis), haemorrhages in the lumen, squamous metaplasia. Histopathological parameters were assessed using a semi-quantitative scoring system ranging from 0 to 4: (0) no lesion; (1) minimal lesion; (2) mild lesion; (3) moderate lesion; (4) severe lesion. The cumulative lesion score for each mouse was calculated, and the group mean score was determined for each experimental condition.

In addition, serial lung sections were used for viral antigen detection by immunohistochemistry. Briefly, sections were deparaffinised and rehydrated through xylene and graded alcohol, respectively, and quenched for endogenous peroxidase activity with 3% hydrogen peroxide in methanol for 30 minutes at room temperature. After that, heat-induced antigen retrieval was achieved submerging the slides in 10mM sodium citrate buffer, pH 3.2, and using a microwave to bring the sections to a sub-boiling temperature for 10 minutes. Slides were then blocked with 10% normal goat serum for 60 minutes, followed by overnight incubation with a rabbit polyclonal primary antibody against vaccinia virus obtained in-house at a 1:500 dilution. Afterward, a secondary antibody (DAKO EnVision labelled polymer goat anti-rabbit) was applied for 1 hour, and antigen expression was visualised using 3,3′-diaminobenzidine tetrahydrochloride (DAB) (Dako REAL EnVision Detection System).

### Virus titration from organs

For determination of virus titres and viral DNA in organs, infected mice were euthanized at 5 dpi, and samples of the indicated organs from each animal were aseptically removed. Samples were weighed, homogenized in 1 ml PBS using Tissue Lyser II (Qiagen), and freeze/thawed 3 times. Then, the amount of infectious virus was determined by plaque assay as described above in BS-C-1 cells. In every case, virus titres were calculated taking into account each organ sample weight after extraction.

### Quantification of MPXV DNA

MPXV DNA from infected right lungs was detected by qPCR using specific oligonucleotides and fluorescent-labelled probes previously described for clade I and II viruses ^30,31^ (see Table S1). DNA was isolated from homogenized MPXV infected organs using Maxwell RSC Viral TNA kit (Promega) and Maxwell RSC instrument (Promega) and finally eluted in 50 ul nuclease free water. We set up a multiplex RT-qPCR to detect viral DNA from both clades together with specific oligonucleotides for the detection of the hypoxanthine phosphoribosyltransferase1 (HPRT1) gene as an extraction control (see supplementary table for nucleotide and probe sequences). 3 ul of isolated DNA or control MPXV DNA were incubated with qPCR mix containing 800 nmol/L forward and reverse primers of both clades, 100 nmol/L forward and reverse primers of HPRT, 200 nmol/L probe with Reliance One-Step Multiplex RT-qPCR Supermix (Bio-RaD). The thermal cycling conditions using a CFX Opus 384 Real-Time PCR System (Bio-Rad) were: one cycle at 50 °C for 10 min, one cycle at 95 °C for 10 min followed by 40 cycles at 95 °C for 10 s and 60 °C for 30 s. Data were analyzed with CFX Maestro 2.3 (Bio-Rad). All measurements were made in triplicate, and a non-template control together with the standard curve were included in each plate (7-log standard curve of 10-fold dilutions of a plasmid containing an 85 bp insert of monkeypox virus DNA for clade IIB and a PCR fragment of 90pb for clade Ib). After quantification by Ct interpolation in the calibration curve, data were converted to copies of MPXV DNA/g.

### Statistical analyses

Statistical analyses were performed using Prism software (GraphPad Software, Inc.). Data from viral growth curves was analyzed using multiple unpaired t tests. For the analysis of survival data, the Log-rank (Mantel-Cox) was used to calculate the P-values. The weight data was analyzed using an unpaired two-tailed Student’s *t* test (Mann-Whitney U-test). The differences in histopathological lesion scores among the experimental groups were determined using an unpaired t-test.

## Results

### MPXV clade Ib shows increased virus spread in cell culture

Previous studies revealed a defect in the replication and spread of the 2022 MPXV compared with its closest ancestor ^26^ and also with other clade II viruses ^24,25,32^, consisting in reduced release of virus to media. Thus, we analyzed the growing kinetics of clade Ib in BS-C-1 and human fibroblasts HFF-1 as compared to 2022 clade IIb. Infection of cells at high multiplicity of infection (moi) rendered similar levels of infectious virus after a single round of infection in the two cell lines examined (Fig. 1a, b). Consistent with this, no significant differences in plaque size between both clades were detected (Fig. 1c). On the contrary, when using low moi, after multiple rounds of infection, we observed higher levels of infectious extracellular virus in the supernatants from clade Ib infected cells compared to those infected with the global clade IIb MPXV (Fig. 1e, g). However, the levels of cell associated virus remained similar between the two clades (Fig. 1d, f). These differences were further confirmed by a comet tail assay in BSC-40 cells, an assay to visualize viral spread through tissue cultures. As previously reported by other groups ^24,26^, the 2022 clade IIb did not induce comet formation whereas the comet phenotype, consequence of the extracellular virus release from infected cells, was clearly evident in clade Ib infected cultures (Fig. 1h). Together, these results indicated more efficient long-range spreading through cell culture for clade Ib compared to the 2022 clade IIb.

**Figure 1.**
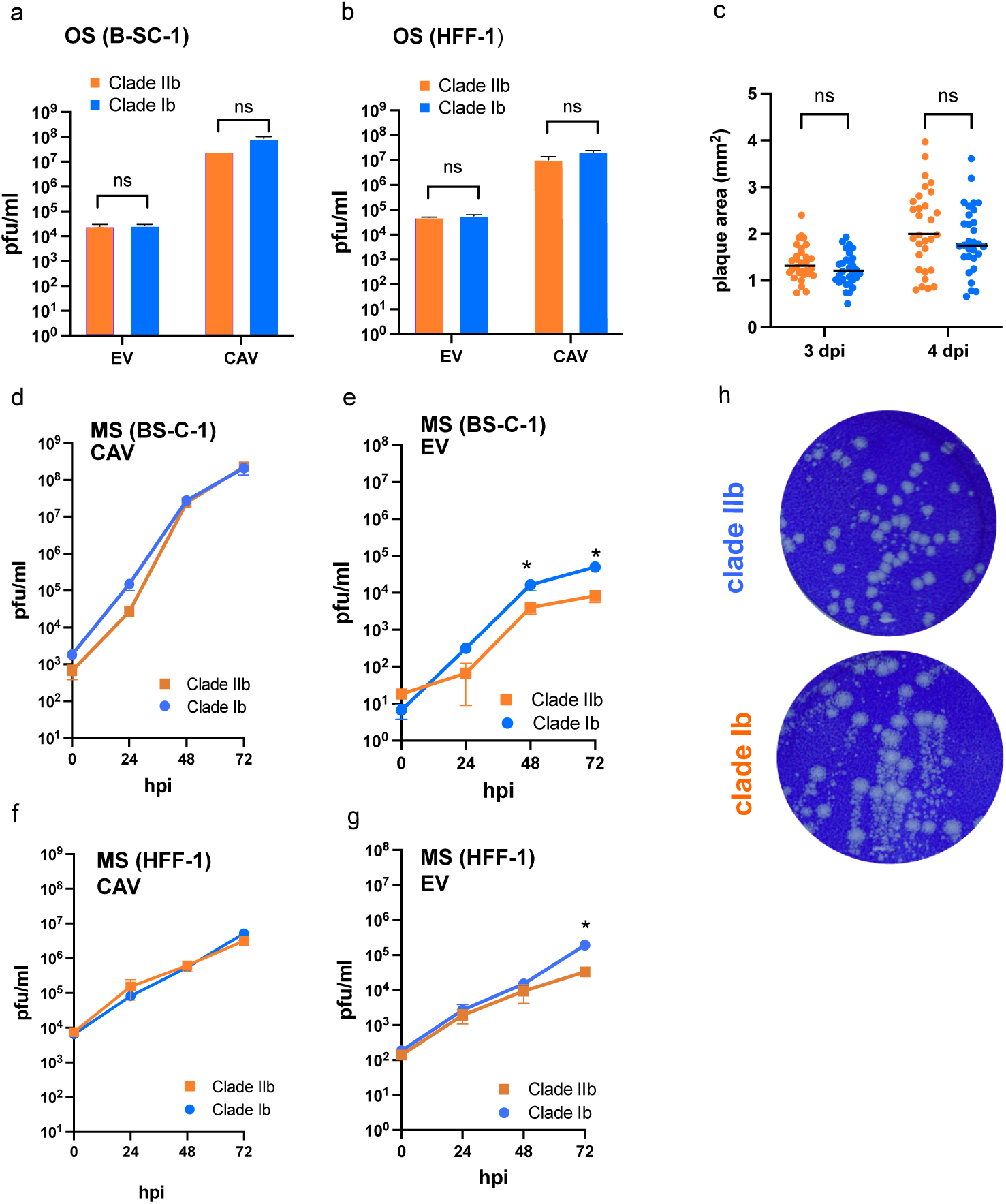
MPXV clade Ib spreads more efficiently than global clade IIb in cell culture. To assess viral replication kinetics of MPXV clades Ib and IIb, BS-C-1 (a, d, e) or HFF-1 (b, f, g) cells were infected with 5 pfu/cell in the one step growth curves (OS, a,b) or 0.05 pfu/cell in the multi-step growth curves (MS, d-g). In every case, cell-associated virus (CAV) and extracellular virus (EV) were determined at the times post infection indicated. Triplicates were used and data shown are mean ± SD. *, p<0.05 using an unpaired t test (c) Viral plaque size quantification at 3 and 4 dpi (n = 30). ns; non-significant. (h) Representative images of the comet assays conducted in BSC-40 with MPXV clades IIb (up) and Ib (down).

### Clade Ib MPXV is more virulent than the global clade IIb in CAST mice

Previous studies have demonstrated, in susceptible CAST mice, the loss of virulence of the clade IIb lineage B.1 MPXV causing the global outbreak in 2022. It is particularly noteworthy the difference in virulence with its immediate ancestor within clade IIb, the A.1 lineage, from which it is separated by only 46 nucleotide changes induced by the action of APOBEC3 as consequence of human-to-human transmission ^26^. The emerging clade Ib has also undergone this type of transmission, especially in Central African regions, and an APOBEC3 signature has also been identified in its genome. We aimed to determine in CAST mice whether this transmission deeply attenuated clade Ib to the levels of the other mpox virus with sustained transmission among humans.

First, we tested in CAST mice the virulence of the MPXV clade IIb from the 2022 global outbreak used in this work, which was isolated in Madrid. Coincident with previous observations from other groups using other clade IIb virus isolates from the 2022 global outbreak ^24,25^, mice intranasally infected with 10^5^ pfu of clade IIb did not developed any signs of disease, weight loss or mortality. Then, we increased the infectious dose up to 2x10^6^ pfu using the same route of infection and a large number of animals, and observed a range of bodyweight loss in the 80% (36/45) of the infected mice (Fig. 2b), with no significant differences between males and females (Fig. 2c, d). Although the average weight loss observed for infected mice was around 10% of the initial bodyweight, in the 20% (9/45) of the inoculated mice this weight loss reached the 20% or higher values (Fig. 2b). The mortality observed was around 9% (4/45) and surviving animals completely recovered from infection by day 11 post infection.

**Figure 2.**
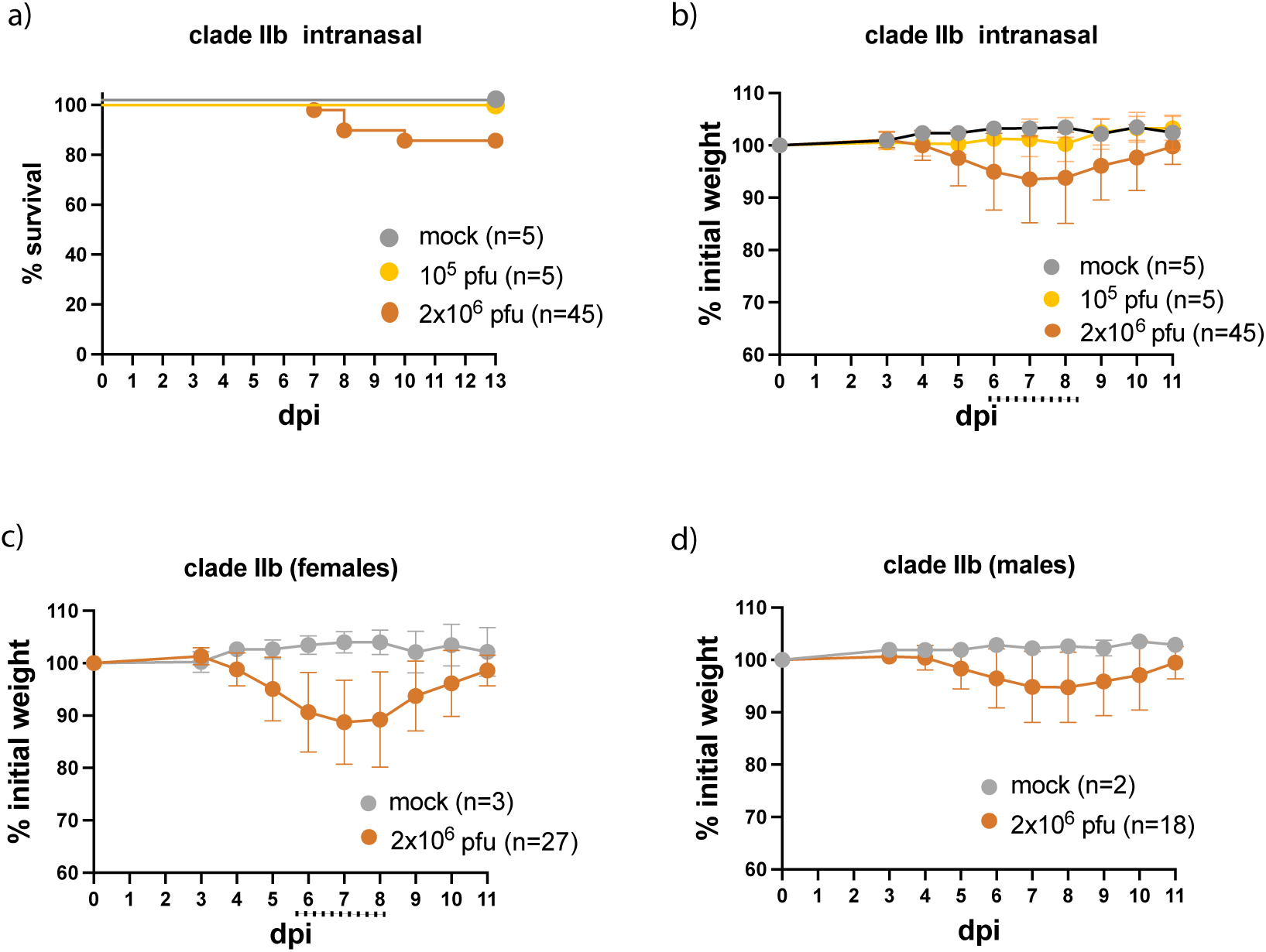
Virulence of MPXV clade IIb upon intranasal infection. CAST mice were mock infected or infected intranasally with 10^5^ or 2 x 10^6^ pfu of MPXV clade IIb lineage B.1 and monitored for survival (a) and bodyweight loss (b). Bodyweight data from mice infected with the highest dose were categorized by sex (c and d). Data correspond to the mean and bar indicates SD and bar indicates SD. Dotted line indicates significant differences in bodyweight (Mann-Whitney U-test, p<0.05) between mock and clade IIb 2 x 10^6^ pfu groups.

In the case of clade Ib, all mice (8/8) infected intranasally with 10^5^ and 10^6^ pfu, as well as 75% (3/4) of those infected with 10^4^ pfu, showed severe weight losses of approximately 25% of their original bodyweight (Fig. 3a) and finally succumbed to the infection within 7-11 days (Fig. 3b). Severe hypothermia was observed in mice infected with 10^5^ and 10^6^ pfu of clade Ib (Fig. 3c). This response has been associated with severe systemic infection and mortality in mice infected with the related orthopoxvirus vaccinia virus ^33^ (Fig. 3c). However, such hypothermia was not observed in mice infected with clade IIb (Fig. 3c), which is consistent with the lack of severe weight loss and mortality in this group (Fig. 3a and b). Similar results were obtained after intraperitoneal MPXV infection to mimic a systemic infection, where all mice (4/4) died within 7 days following infection with 10^5^ pfu of clade Ib (Fig. 3d, e). In this case, the weight losses observed were less pronounced compared to the intranasal inoculation. However, no mortality or weight loss was observed when clade IIb was administered intraperitoneally at the same dosage (Fig. 3d, e).

**Figure 3.**
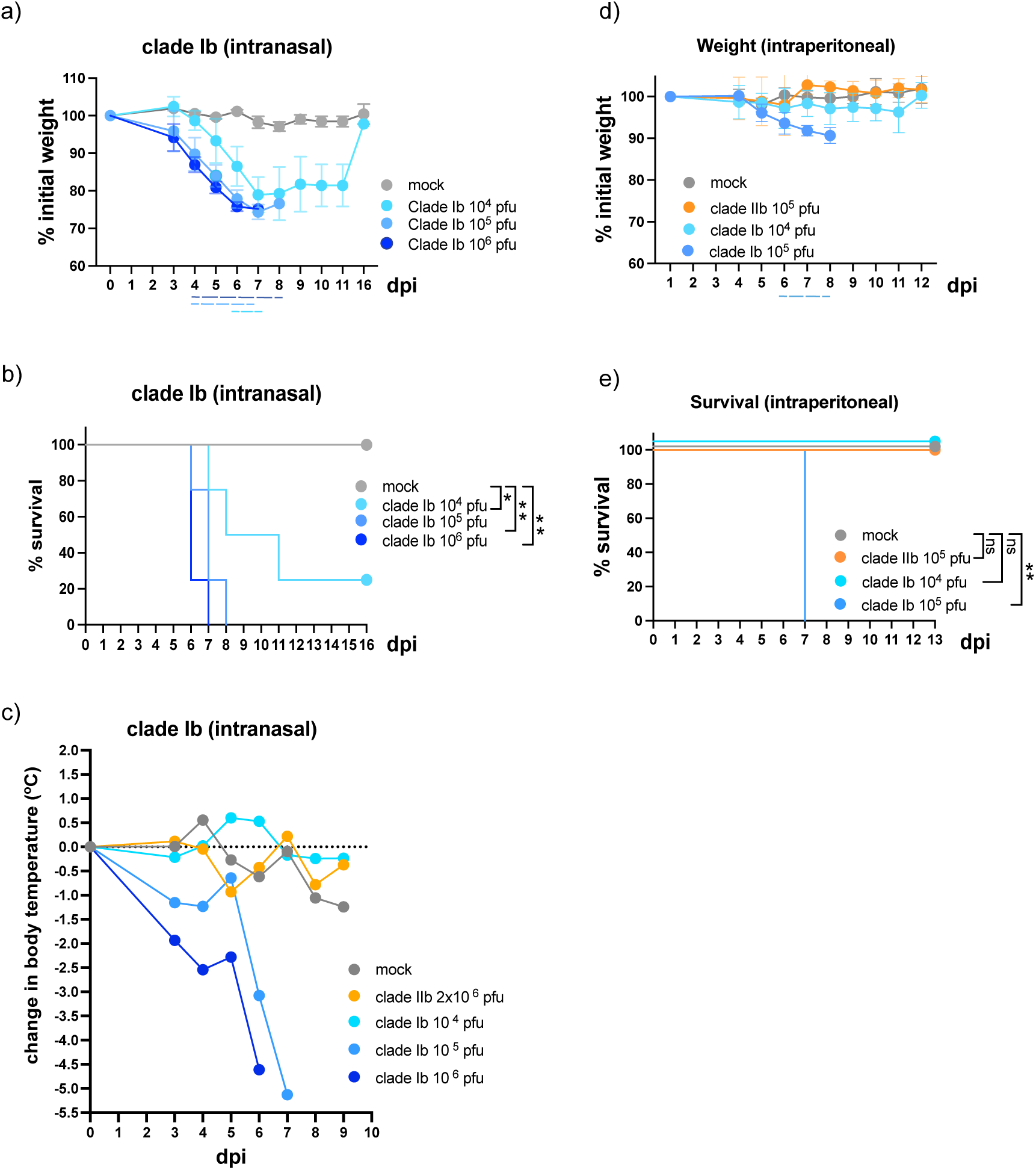
MPXV clade Ib is more virulent than clade IIb. Groups of CAST mice (n=4) were intranasally (a - c) mock infected or infected with increasing doses of MPXV clade Ib and monitored daily for changes in bodyweight (a), survival (b) and body temperature (c). For comparison, body temperature data from mice intranasally infected with 2 x 10^6^ pfu of clade IIb (n=4) are also shown in panel (c), while weight and survival data for this group are excluded from panels (a) and (b) to enhance clarity, as no severe weight loss or mortality were detected. Similarly, groups of mice (n=4) were daily monitored after intraperitoneal infection (d, e) with the indicated doses of clades Ib and IIb. Data correspond to the mean and bar indicates SD. ns, no significant; *, p<0.05; **, p<0.01 in Survival Log-rank (Mantel-Cox) test and discontinuous line indicates significant differences in bodyweight (Mann-Whitney U-test, p<0.05) between mock and the corresponding group indicated by colour legend.

### MPXV clade Ib intranasal infection results in more severe lung pathology than global clade IIb

We performed histopathological evaluations of lung tissue sections from infected mice at 5 dpi to examine the pathology caused by both MPXV clades following intranasal infection with different doses. Overall, lung lesions were more severe and widespread in CAST mice infected with clade Ib compared to those infected with the global clade IIb lineage B.1. Although the difference in histopathological scores did not reach statistical significance, the trend toward higher scores in the clade Ib supports its greater virulence observed (Fig. 4a and b). The lesions observed after intranasal infection with 10^5^ pfu of both MPXV clades included moderate septal thickening (interstitial pneumonia), small alveolar haemorrhages, alveolar cellular infiltrates consisting mainly of macrophages and occasional lymphocytes, bronchiolar epithelial hyperplasia, bronchioles with epithelial necrosis, sloughed epithelium and inflammatory mononuclear infiltrates (bronchiolitis), as well as moderate perivascular and peribronchiolar mononuclear infiltrates along with occasional degenerative and viable neutrophils (Fig. 4c). In mice infected with the same dose of clade Ib, the lesions were similar but more severe and extensive, also presenting moderate to severe perivascular oedema, haemorrhages in the bronchiolar lumen, necrosis and desquamation of alveolar septa, and abundant perivascular and peribronchiolar degenerative and viable neutrophils (Fig. 4c). In mice infected with a higher dose (10^6^ pfu) of both clades, the lesions were similar but more severe and diffuse that those described in mice infected with 10^5^ pfu. Once again, tissue destruction - particularly necrosis and desquamation of alveolar septa - and vascular changes such as perivascular and alveolar oedema were more pronounced in mice infected with clade Ib (Fig. 4c).

**Figure 4.**
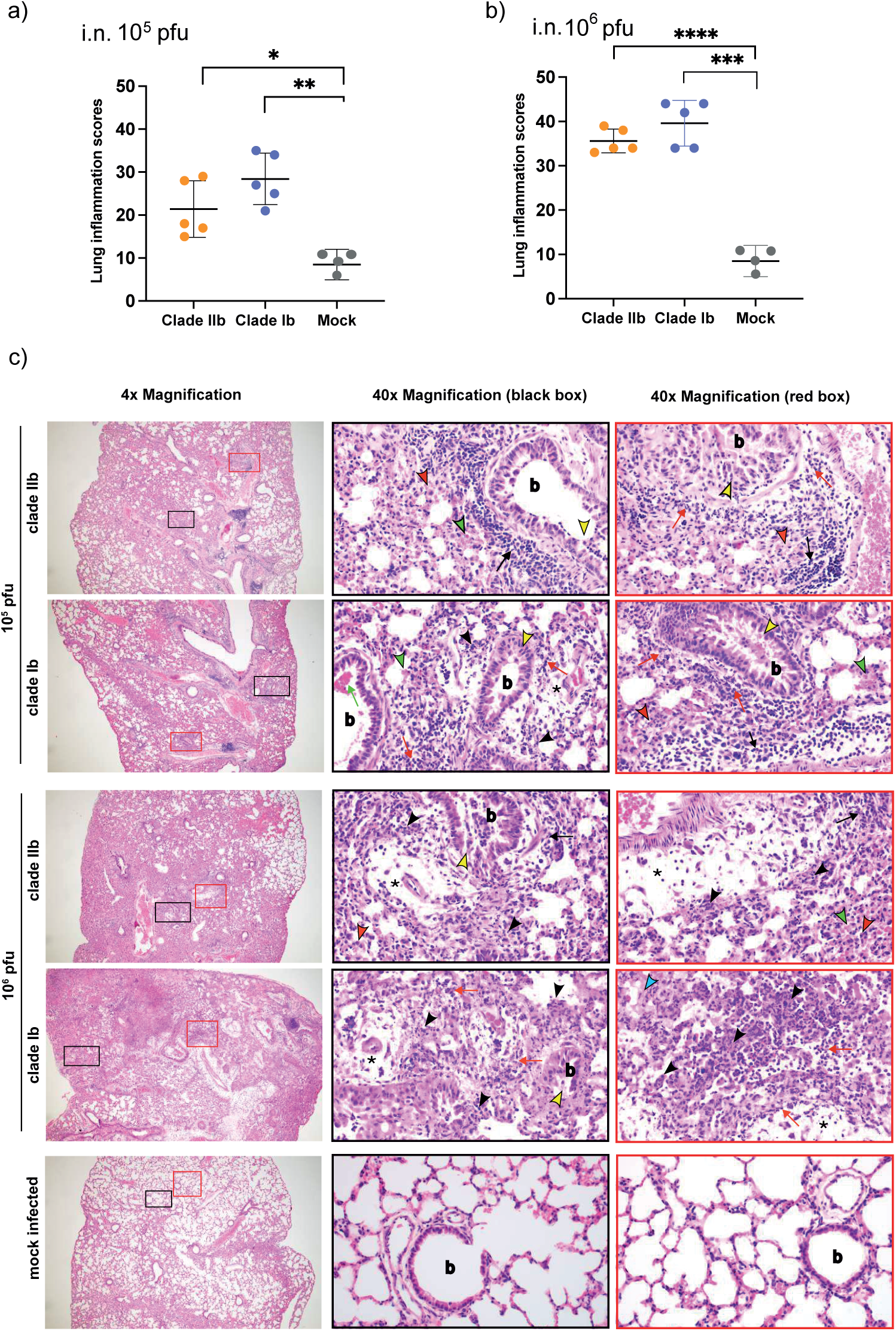
Comparative lung pathology of clades IIb and Ib after intranasal infection. Groups of CAST mice (n = 5) were intranasally infected with 10^5^ or 10^6^ pfu of clades IIb and Ib and lung tissue examined at 5 dpi. (a-b) Mean and SD of cumulative histopathological lesion scores in lung samples from mock and intranasally infected mice. *p < 0.05; **p < 0.01; ***p < 0.001; ****p < 0.0001. (c) Representative histopathological lung sections (haematoxylin and eosin staining). A general view of the lung area along with histopathological details from selected lung areas (black and red boxes) are shown. Note the absence of significant lesions in mock-infected mice, where alveolar spaces without inflammatory changes and empty airways without airway epithelial involvement predominate compared to mice infected at different doses. Legend: b, bronchioles; green arrowheads, small alveolar haemorrhages; red arrowheads, alveolar cellular infiltrates mainly of macrophages and occasional lymphocytes; yellow arrowheads, bronchiolitis; black arrows, moderate perivascular and peribronchiolar mononuclear infiltrates; red arrows, occasional degenerative and viable neutrophils; black asterisks, severe perivascular oedema; green arrows, haemorrhages in the bronchiolar lumen; black arrowheads, necrosis and desquamation of alveolar septa; red arrows, abundant perivascular and peribronchiolar degenerative and viable neutrophils; blue arrowheads, perivascular and alveolar oedema.

Immunohistochemical detection of MPXV in the lungs of infected mice (Fig. 5) demonstrated that viral antigen from both clades was predominantly localized in the epithelial cells of the bronchioles and in adjacent lung tissue, particularly in the pneumocytes and the interstitial macrophages within the thickened alveolar septa. Viral antigen was also observed in macrophages infiltrating bronchiolar epithelium, mononuclear cells in the lumen of the bronchioles, endothelial cells and interstitial fusiform-shaped cells, identified mainly as fibroblasts and occasionally smooth muscle cells. However, a higher number of lung regions positive for viral antigen, overlapping with areas of tissue damage, were observed in those mice infected with clade Ib. This finding aligns with the increased severity and extent of lung lesions induced by clade Ib infection (Fig. 5).

**Figure 5.**
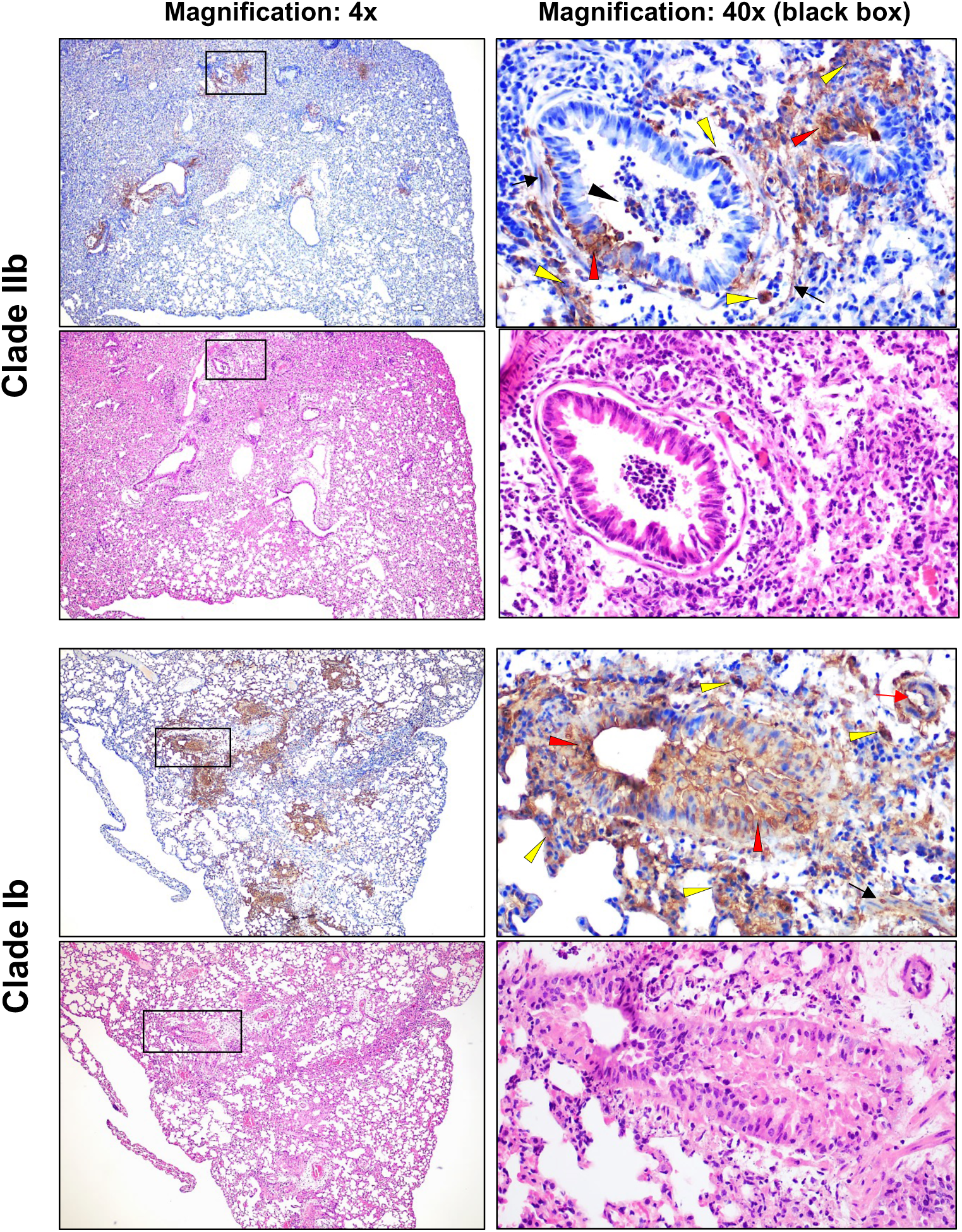
MPXV distribution in the infected lung. Representative images of serial lung sections were taken from Cast/EiJ mice that were intranasally infected with 10⁶ pfu of MPXV clades Ib and IIb. The sections were immunolabelled against MPXV, or stained with haematoxylin and eosin for histopathological evaluations. A general view of the lung areas is shown (magnification: 4x), alongside details of selected lung areas (black boxes, magnification: 40x). Note the higher number of lung areas positive for viral antigen with clade Ib, which was in correlation with the presence of greater tissue damage Cell types and areas positive for viral antigen: red arrowheads, epithelial cells of the bronchioles, yellow arrowheads, pneumocytes and interstitial macrophages in the thickened alveolar septa; red arrows, endothelial cells; black arrowheads, macrophages infiltrating bronchiolar epithelium and mononuclear cells in the lumen of the bronchioles; black arrows, fibroblasts and smooth muscle cells.

### MPXV clade Ib disseminates to internal organs more efficiently than clade IIb

The previously observed differences in extracellular virus production (Fig. 1), between clade Ib and clade IIb lineage B.1 *in vitro* (Fig. 1) may influence viral dissemination within the host following infection and could partly account for the virulence differences observed between these two MPXV clades (Fig. 2 and 3). To address this, we determined viral loads in various organs five days after intranasal infection of CAST mice with either 10^5^ or 10^6^ pfu. High viral titers were detected in the nasal turbinates, for both MPXV clades, indicating efficient replication at the site of inoculation. Notably, significant differences between clades in nasal turbinates were apparent only at the lower infectious dose. In the lungs, clade Ib consistently exhibited a significant higher level of infectious virus than clade IIb across both inoculation doses (Fig. 6a, b). Importantly, infectious virus was detected in the spleens of mice infected with clade Ib, but not clade IIb, at either dose (Fig. 6a, b), indicating systemic dissemination to internal organs. These findings were confirmed by determination of viral genome copies in lung and spleen tissues (Fig. 6c-e). Overall, the data show that clade Ib replicates more efficiently in the lungs and is capable of reaching internal organs, contributing to increased lethality in infected mice.

**Figure 6.**
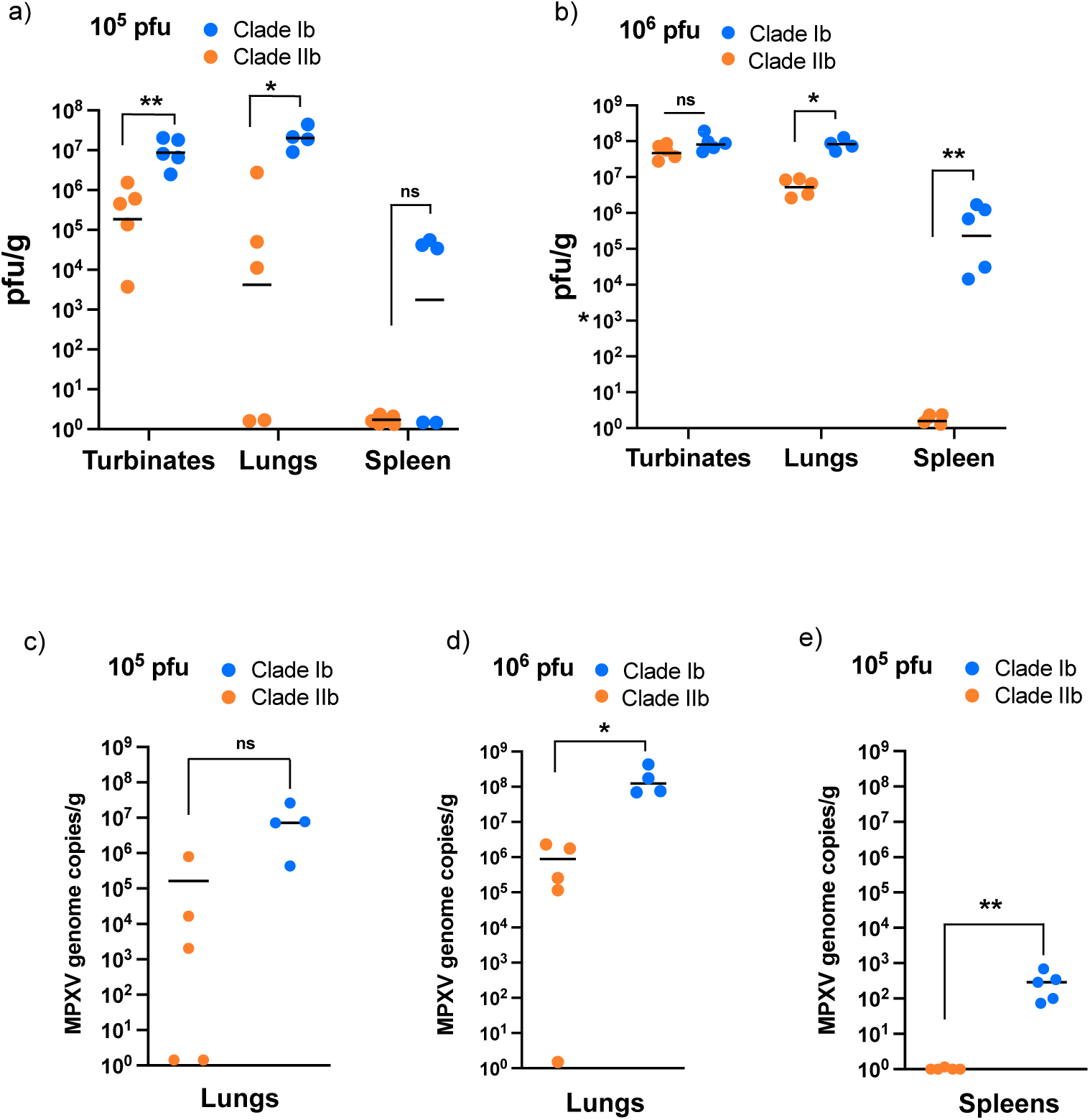
Viral loads in MPXV clade Ib disseminates to internal organs more efficiently than clade IIb. CAST mice were intranasally infected with either 10^5^ (a, c, e) or 10^6^ (b, d) pfu of MPXV clades Ib and IIb. Five days after infection infectious virus (a, b) and MPXV genome copies (c - e) were determined from the indicated organs. Dots in all panels represent individual sample data and bar indicates geometric mean of the group (n=5, except for clade Ib lungs). *, p<0.05, **, p<0.01 (Mann-Whitney U-test).

## Discussion

Since May 2022, two Public Health Emergencies of International Concern have been declared by WHO caused by mpox outbreaks characterized by sustained human-to-human transmission. During the first one, the 2022 global mpox outbreak, emerging clade IIb lineage B.1 spread predominantly through sexual networks particularly among men who have sex with men (MSM), and was associated with a crude death rate of 0.1 % ^34^. The recent second Public Health Emergency of International Concern, in 2024, involved the novel clade Ib which has rapidly disseminated among humans across Central African regions via both sexual and non-sexual routes of transmission, and although its associated case fatality rate was estimated to be higher than clade IIb, there are still some concerns related to its severity ^13,14,35^. Despite both clade IIb and clade Ib displaying clear APOBEC3 mutational signatures associated with sustained human transmission ^12,36^, our data indicate different biological behaviour for these lineages. While clade IIb lineage B.1 exhibits reduced viral spread *in vitro*, clade Ib produces significantly higher extracellular virus and displays enhanced cell-to-cell dissemination in cell culture.

Previous studies using the CAST mice model of MPXV infection, which has been proposed as a feasible murine model of MPXV infection to compare the virulence of the diverse existing clades, showed that clade IIb lineage B.1 is hugely attenuated compared to its antecessor clade IIb lineage A.1 ^25,26^ and previous clade IIa lineages ^24,25^. The data presented here, using this animal model, show that the recently emerged clade Ib is considerably more virulent than the global clade IIb from 2022, causing higher mortality with lower viral doses, more severe lung pathology and successful dissemination to internal organs. All mice succumbed to infection after intranasal or intraperitoneal inoculation of 10^5^ pfu, similar to clade IIa USA-2003 isolate but far from the mortalities observed with reference clade I viruses like Zaire-1979 isolate, according to previous data from *Americo et al*. This would mean that clade Ib is not as virulent as previous clade I viruses in CAST mice and might have been somehow attenuated after sustained human-to-human transmission, since Zaire-1979 likely originated from a single spillover from an unknown animal reservoir with limited human-to-human transmission ^37^. These virulence differences could be attributed to distinct genomic changes accumulated in clade Ib relative to ancestral clade I viruses. Recent genomic analyses have identified several mutations in genes critical for viral replication (such as L6R/OPG105 and A25R/OPG151) and viral immunomodulation in the clade 1b viruses. Interestingly, C9L (OPG047), the gene encoding the only Kelch-like protein in MPXV and involved in immune evasion, is affected by an in-frame deletion and three aminoacid substitutions. Another immunomodulatory gene previously associated to MPXV virulence ^38^ and affected by a large deletion in clade Ib is D14L (OPG032), which encodes the MPXV complement control protein and is also absent from clade II viruses ^3,4^. Also, the large and immunogenic surface glycoprotein B21R (OPG210) accumulates diverse aminoacid substitutions that might affect its ability to suppress T-cell responses.

Importantly, although the CAST mice model of MPXV infection using intranasal or intraperitoneal inoculations does not fully recapitulate the clinical features of human mpox, it correlates accurately with the relative severity of the diverse clades in humans ^24^, as diminished virulence for mice of clade IIb correlates with its transmission in humans during the 2022 global outbreak. Moreover, there is increasing evidence indicating that mpox will remain a public health threat in the near future, affecting both endemic and non-endemic regions ^20,39^. The current epidemiological context, characterized by sustained human-to-human transmission, facilitates APOBEC3-driven evolution of MPXV within highly virulent clades, raising the possibility of emergent viral lineages with unpredictable pathogenic potential. Therefore, the data presented here using the CAST mouse model have important implications for guiding future public health preparedness and surveillance strategies against mpox.

In summary, our findings indicate that the recently emerged clade Ib MPXV retains a higher level of virulence compared to the 2022 global outbreak clade IIb. Continuous genomic and functional surveillance of newly arising MPXV lineages will anticipate shifts in viral virulence and transmissibility, particularly as sustained human-to-human transmission continues in both endemic and non-endemic regions.

## Acknowledgements

This work was supported by grants PID2022-136867NB-I00 and RTI2021-128580OB-100 from the Spanish Ministry of Science, Innovation and Universities to BH and AA, respectively and grant HR23-00558 from La Caixa Foundation to AA. ARU was supported by a PhD studentship (PRE2022-103462) from the Spanish Ministry of Science, Innovation and Universities.

## Author contributions

Conceptualization: AA and BH. Design and supervision of the experiments: BH. Experimentation: ARU, BH, CS, LFA, PJS and RS. Formal analysis: AA, ARU, BH, PJS. Writing the original draft of the manuscript: BH. Review of the original draft: AA, BH, PJS. Funding acquisition: AA and BH. All authors revised and agreed to the final version of the manuscript.

