## Supplemental Tables S1 and S2 for "Enhanced mouse virulence of mpox virus clade Ib over clade IIb despite genomic changes caused by human-to-human transmission"

### Supplementary Material

#### Table of contents

#### Supplementary Tables

Tables S1 – S2.

**Table S1. CAST EiJ mice from experiments in Figure 2 and 3.** Details about the mice used for each of the experiments in figures from the main text are provided in A-C. n/a: not aplicable.

| **A. Mice from experiment in Figure 2.** | | | | |
|---|---|---|---|---|
| **Group** | **Dose (pfu)** | **Route** | **Age (weeks)** | **Sex** |
| Mock (PBS) | n/a | IN | 8 | Male |
| Mock (PBS) | n/a | IN | 8 | Male |
| Mock (PBS) | n/a | IN | 8 | Female |
| Mock (PBS) | n/a | IN | 8 | Female |
| Mock (PBS) | n/a | IN | 8 | Female |
| Clade IIb | $10^5$ | IN | 7 | Female |
| Clade IIb | $10^5$ | IN | 7 | Female |
| Clade IIb | $10^5$ | IN | 8 | Female |
| Clade IIb | $10^5$ | IN | 8 | Female |
| Clade IIb | $10^5$ | IN | 8 | Female |
| Clade IIb | $2x10^6$ | IN | 7 | Male |
| Clade IIb | $2x10^6$ | IN | 7 | Male |
| Clade IIb | $2x10^6$ | IN | 8 | Male |
| Clade IIb | $2x10^6$ | IN | 8 | Male |
| Clade IIb | $2x10^6$ | IN | 8 | Male |
| Clade IIb | $2x10^6$ | IN | 8 | Female |
| Clade IIb | $2x10^6$ | IN | 8 | Female |
| Clade IIb | $2x10^6$ | IN | 8 | Female |
| Clade IIb | $2x10^6$ | IN | 8 | Female |
| Clade IIb | $2x10^6$ | IN | 8 | Female |
| Clade IIb | $2x10^6$ | IN | 8 | Female |
| Clade IIb | $2x10^6$ | IN | 8 | Female |
| Clade IIb | $2x10^6$ | IN | 8 | Male |
| Clade IIb | $2x10^6$ | IN | 8 | Male |
| Clade IIb | $2x10^6$ | IN | 8 | Male |
| Clade IIb | $2x10^6$ | IN | 8 | Male |
| Clade IIb | $2x10^6$ | IN | 8 | Male |
| Clade IIb | $2x10^6$ | IN | 9 | Female |
| Clade IIb | $2x10^6$ | IN | 8 | Female |
| Clade IIb | $2x10^6$ | IN | 8 | Female |
| Clade IIb | $2x10^6$ | IN | 9 | Female |
| Clade IIb | $2x10^6$ | IN | 9 | Male |
| Clade IIb | $2x10^6$ | IN | 9 | Male |
| Clade IIb | $2x10^6$ | IN | 9 | Male |
| Clade IIb | $2x10^6$ | IN | 8 | Male |
| Clade IIb | $2x10^6$ | IN | 8 | Male |
| Clade IIb | $2x10^6$ | IN | 8 | Female |
| Clade IIb | $2x10^6$ | IN | 8 | Female |
| Clade IIb | $2x10^6$ | IN | 8 | Female |
| Clade IIb | $2x10^6$ | IN | 8 | Female |

| | | | | |
|---|---|---|---|---|
| Clade IIb | 2x10$^6$ | IN | 12 | Female |
| Clade IIb | 2x10$^6$ | IN | 12 | Female |
| Clade IIb | 2x10$^6$ | IN | 11 | Female |
| Clade IIb | 2x10$^6$ | IN | 11 | Female |
| Clade IIb | 2x10$^6$ | IN | 12 | Female |
| Clade IIb | 2x10$^6$ | IN | 12 | Female |
| Clade IIb | 2x10$^6$ | IN | 12 | Female |
| Clade IIb | 2x10$^6$ | IN | 12 | Female |
| Clade IIb | 2x10$^6$ | IN | 12 | Female |
| Clade IIb | 2x10$^6$ | IN | 12 | Female |
| Clade IIb | 2x10$^6$ | IN | 12 | Female |
| Clade IIb | 2x10$^6$ | IN | 12 | Female |
| Clade IIb | 2x10$^6$ | IN | 12 | Female |
| Clade IIb | 2x10$^6$ | IN | 12 | Female |
| Clade IIb | 2x10$^6$ | IN | 12 | Female |

**B. Mice from experiment in Figure 3 a-c.**

| Group | Dose (pfu) | Route | Age (weeks) | Sex |
|---|---|---|---|---|
| Mock (PBS) | n/a | IN | 11 | Female |
| Mock (PBS) | n/a | IN | 11 | Female |
| Mock (PBS) | n/a | IN | 9 | Female |
| Mock (PBS) | n/a | IN | 9 | Female |
| Clade IIb | 2x10$^6$ | IN | 8 | Female |
| Clade IIb | 2x10$^6$ | IN | 8 | Female |
| Clade IIb | 2x10$^6$ | IN | 7 | Female |
| Clade IIb | 2x10$^6$ | IN | 7 | Female |
| Clade Ib | 10$^4$ | IN | 7 | Female |
| Clade Ib | 10$^4$ | IN | 7 | Female |
| Clade Ib | 10$^4$ | IN | 7 | Female |
| Clade Ib | 10$^4$ | IN | 7 | Female |
| Clade Ib | 10$^5$ | IN | 7 | Female |
| Clade Ib | 10$^5$ | IN | 7 | Female |
| Clade Ib | 10$^5$ | IN | 7 | Female |
| Clade Ib | 10$^5$ | IN | 7 | Female |
| Clade Ib | 10$^6$ | IN | 7 | Female |
| Clade Ib | 10$^6$ | IN | 7 | Female |
| Clade Ib | 10$^6$ | IN | 7 | Female |
| Clade Ib | 10$^6$ | IN | 7 | Female |

| C. Groups of CAST EiJ mice from experiment in Figure 3 d-e. | | | | |
|---|---|---|---|---|
| **Group** | **Dose (pfu)** | **Route** | **Age (weeks)** | **Sex** |
| Mock (PBS) | n/a | IP | 8 | Male |
| Mock (PBS) | n/a | IP | 8 | Male |
| Mock (PBS) | n/a | IP | 8 | Female |
| Mock (PBS) | n/a | IP | 8 | Female |
| Clade IIb | $10^5$ | IP | 9 | Male |
| Clade IIb | $10^5$ | IP | 9 | Male |
| Clade IIb | $10^5$ | IP | 9 | Female |
| Clade IIb | $10^5$ | IP | 9 | Female |
| Clade Ib | $10^3$ | IP | 8 | Male |
| Clade Ib | $10^3$ | IP | 8 | Female |
| Clade Ib | $10^3$ | IP | 8 | Female |
| Clade Ib | $10^3$ | IP | 8 | Female |
| Clade Ib | $10^4$ | IP | 9 | Female |
| Clade Ib | $10^4$ | IP | 9 | Male |
| Clade Ib | $10^4$ | IP | 9 | Male |
| Clade Ib | $10^4$ | IP | 9 | Male |
| Clade Ib | $10^5$ | IP | 8 | Male |
| Clade Ib | $10^5$ | IP | 9 | Male |
| Clade Ib | $10^6$ | IP | 9 | Male |
| Clade Ib | $10^6$ | IP | 9 | Male |

**Table S2**. Oligonucleotides and fluorescent-labelled probes used in the study.

| Name | Sequence (5'-3') | Description | Ref. |
|---|---|---|---|
| MPXV_WA_fw | CACACCGTCTCTTCCACAGA | Clade II | Li *et al,* 2010 |
| MPXV_WA_rv | GATACAGGTTAATTTCCACATCG | Clade II | Li *et al,* 2010 |
| MPXV WA-probe | 6-FAM-AACCCGTCGTAACCAGCAATACATTT-BHQ-1 | Clade II | Li *et al,* 2010 |
| HPRT1 fw | TTCTTTGCTGACCTGCTGGA | HPRT gene | own design |
| HPRT1_probe | /5HEX/ATTGGTGGA/ZEN/GATGATCTCTCAACT/3IABkFQ/ | HPRT gene | own design |
| HPRT1_rv | ACAATCAAGACATTCTTTCCAGTT | HPRT gene | own design |
| MPOX-C1b-FW | AAGACTTCCAAACTTAATCACTCCT | Clade Ib | Schuele *et al,* 2024 |
| MPOX-C1b-RV | CGTTTGATATAGGATGTGGACATTT | Clade Ib | Schuele *et al,* 2024 |
| MPOX-C1b-probe | /5Cy5/ATATTCAGG/TAO/CGCATATCCACCCACGT/3IAbRQSp/FA | Clade Ib | Schuele *et al,* 2024 |
